# FROG Analysis Ensures the Reproducibility of Genome Scale Metabolic Models

**DOI:** 10.1101/2024.09.24.614797

**Authors:** Karthik Raman, Miroslav Kratochvíl, Brett G. Olivier, Matthias König, Pratyay Sengupta, Dinesh Kumar Kuppa Baskaran, Tung V N Nguyen, Daniel Lobo, St Elmo Wilken, Krishna Kumar Tiwari, Aswathy K. Raghu, Indumathi Palanikumar, Lavanya Raajaraam, Maziya Ibrahim, Sanjaay Balakrishnan, Shreyansh Umale, Frank Bergmann, Tanisha Malpani, Venkata P Satagopam, Reinhard Schneider, Moritz E. Beber, Sarah Keating, Mihail Anton, Alina Renz, Meiyappan Lakshmanan, Dong-Yup Lee, Lokanand Koduru, Reihaneh Mostolizadeh, Oscar Dias, Emanuel Cunha, Alexandre Oliveira, Yi Qing Lee, Karsten Zengler, Rodrigo Santibáñez-Palominos, Manish Kumar, Matteo Barberis, Bhanwar Lal Puniya, Tomáš Helikar, Hoang V. Dinh, Patrick F. Suthers, Costas D. Maranas, Isabella Casini, Seyed Babak Loghmani, Nadine Veith, Nantia Leonidou, Feiran Li, Yu Chen, Jens Nielsen, GaRyoung Lee, Sang Mi Lee, Gi Bae Kim, Pedro T. Monteiro, Miguel C. Teixeira, Hyun Uk Kim, Sang Yup Lee, Ulf W. Liebal, Lars M. Blank, Christian Lieven, Chaimaa Tarzi, Claudio Angione, Manga Enuh Blaise, Çelik Pınar Aytar, Mikhail Kulyashov, llya Akberdin, Dohyeon Kim, Sung Ho Yoon, Zhaohui Xu, Jyotshana Gautam, William T. Scott, Peter J. Schaap, Jasper J. Koehorst, Cristal Zuñiga, Gabriela Canto-Encalada, Sara Benito-Vaquerizo, Ivette Parera Olm, Maria Suarez-Diez, Qianqian Yuan, Hongwu Ma, Mohammad Mazharul Islam, Jason A. Papin, Francisco Zorrilla, Kiran Raosaheb Patil, Arianna Basile, Juan Nogales, Granado San León, Freddy Castillo-Alfonso, Roberto Olivares-Hernández, Gabriela Canto-Encalada, Gabriel Vigueras-Ramírez, Henning Hermjakob, Andreas Dräger, Rahuman S Malik-Sheriff

## Abstract

Genome-scale metabolic models (GEMs) and other constraint-based models (CBMs) play a pivotal role in understanding biological phenotypes and advancing research in areas like metabolic engineering, human disease modelling, drug discovery, and personalized medicine. Despite their growing application, a significant challenge remains in ensuring the reproducibility of GEMs, primarily due to inconsistent reporting and inadequate model documentation of model results. Addressing this gap, we introduce FROG analysis, a community-driven initiative aimed at standardizing reproducibility assessments of CBMs and GEMs. The FROG framework encompasses four key analyses—Flux variability, Reaction deletion, Objective function, and Gene deletion—to produce standardized, numerically reproducible FROG reports. These reports serve as reference datasets, enabling model evaluators, curators, and independent researchers to verify the reproducibility of GEMs systematically.

BioModels, a leading repository of systems biology models, has integrated FROG analysis into its curation workflow, enhancing the reproducibility and reusability of submitted GEMs. In our study evaluating 65 GEM submissions from the community, approximately 40% reproduced without intervention, 28% requiring minor adjustments, and 32% needing input from authors. The standardization introduced by FROG analysis facilitated the detection and resolution of issues, ultimately leading to the successful reproduction of all models. By establishing a standardized and comprehensive approach to evaluating GEM reproducibility, FROG analysis significantly contributes to making CBMs and GEMs more transparent, reusable, and reliable for the broader scientific community.

## Main Article

Genome-scale metabolic models (GEMs) - the constraint-based models (CBMs) generated from a genomic reconstruction of metabolic pathways - are pivotal in the study of biological phenotypes (Schellenberger et al., 2011). GEMs and other CBMs have broad applications, ranging from understanding microbial, plant, and mammalian metabolism to producing chemicals and materials through metabolic engineering (McCloskey et al., 2013; Oberhardt et al., 2009). They can also be applied to predict enzyme functions and study host-pathogen interactions, microbial interactions in communities (Ibrahim et al., 2021), and cell-cell interactions (van der Ark et al., 2017). Recently, GEMs have been used to advance our understanding of human diseases, and the scope of GEMs has expanded to include drug discovery and personalised medicine (Li et al., 2023; Renz et al., 2020). The CBMs and GEMs have evolved over the past four decades as one of the prominent systems biology modelling approaches, with an increasing number of studies combining models with high-throughput data for efficient predictions (Gu et al., 2019).

However, a substantial challenge with these models is their reproducibility - the ability to reproduce the published results - which is often due to insufficient or inconsistent reporting of model parameters, constraints, and quantitative predictions (Ravikrishnan and Raman, 2015). Metabolic flux values from commonly reported flux balance analyses of CBMs are not unique solutions and do not suffice for reproducibility assessment. Often, the inadequate reporting of objective functions further thwarts verifying whether the publicly shared model aligns with the one used in the study, casting doubt on the scientific validity.

A study (Tiwari et al., 2021) highlighted that approximately half of the selected ordinary differential equation (ODE) models published in peer-reviewed journals could not be reproduced using the information provided in the publications. GEMs are also anticipated to face a comparable crisis in reproducibility. About 9% of the ODE models could be empirically corrected and reproduced through a trial-and-error approach. Such an approach to correct GEMs is impractical, as they are often very large models encompassing thousands of reactions and parameters. To address this, the metabolic model test suite MEMOTE was developed as a standardised framework to assess GEM quality regarding stoichiometry, mass balance, and annotation (Lieven et al., 2020). There are also efforts to standardise GEMs reconstruction (Anton et al., 2023). However, these initiatives didn’t address the reproducibility of the model simulations, urging the scientific community to build upon these foundational efforts.

To address this challenge, we initiated a community effort to standardise the assessment of model reproducibility by developing a new framework - FROG analysis. FROG is an ensemble of analyses for constraint-based models that generate standardised, numerically reproducible reference datasets, termed ‘FROG Reports’. FROG encompasses (1) **F**lux variability, (2) **R**eaction deletion, (3) **O**bjective function, and (4) **G**ene deletion analyses (Figure 1). A FROG report includes four standardised files: (1) upper and lower flux bounds calculated from the flux variability analysis; (2) the vector of objective function values after systematic one-at-a-time reaction deletion, (3) Objective function value of the optimised CBMs, and (4) objective function values vector obtained after each single-gene deletion analysis. Our community recommendation includes public sharing of these reports alongside CBMs and GEMs to enable verification that the same results can be reproduced using the model.

**Figure 1:**
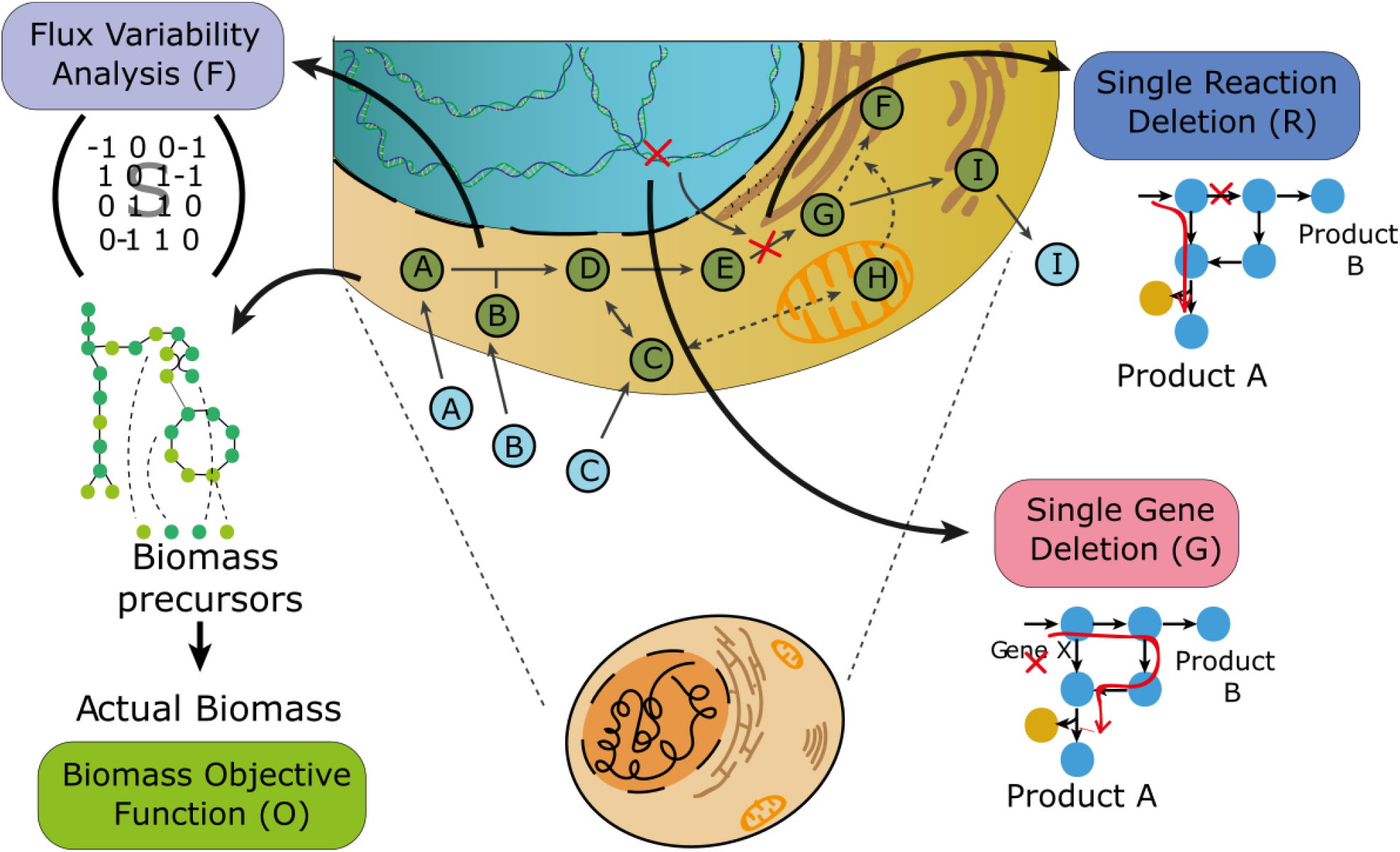
Graphical overview of FROG analysis encompassing (1) Flux variability, (2) Reaction deletion, (3) Objective function, and (4) Gene deletion analyses enable the generation of numerically reproducible reference datasets to assess the reproducibility of GEMs.

**Figure 2:**
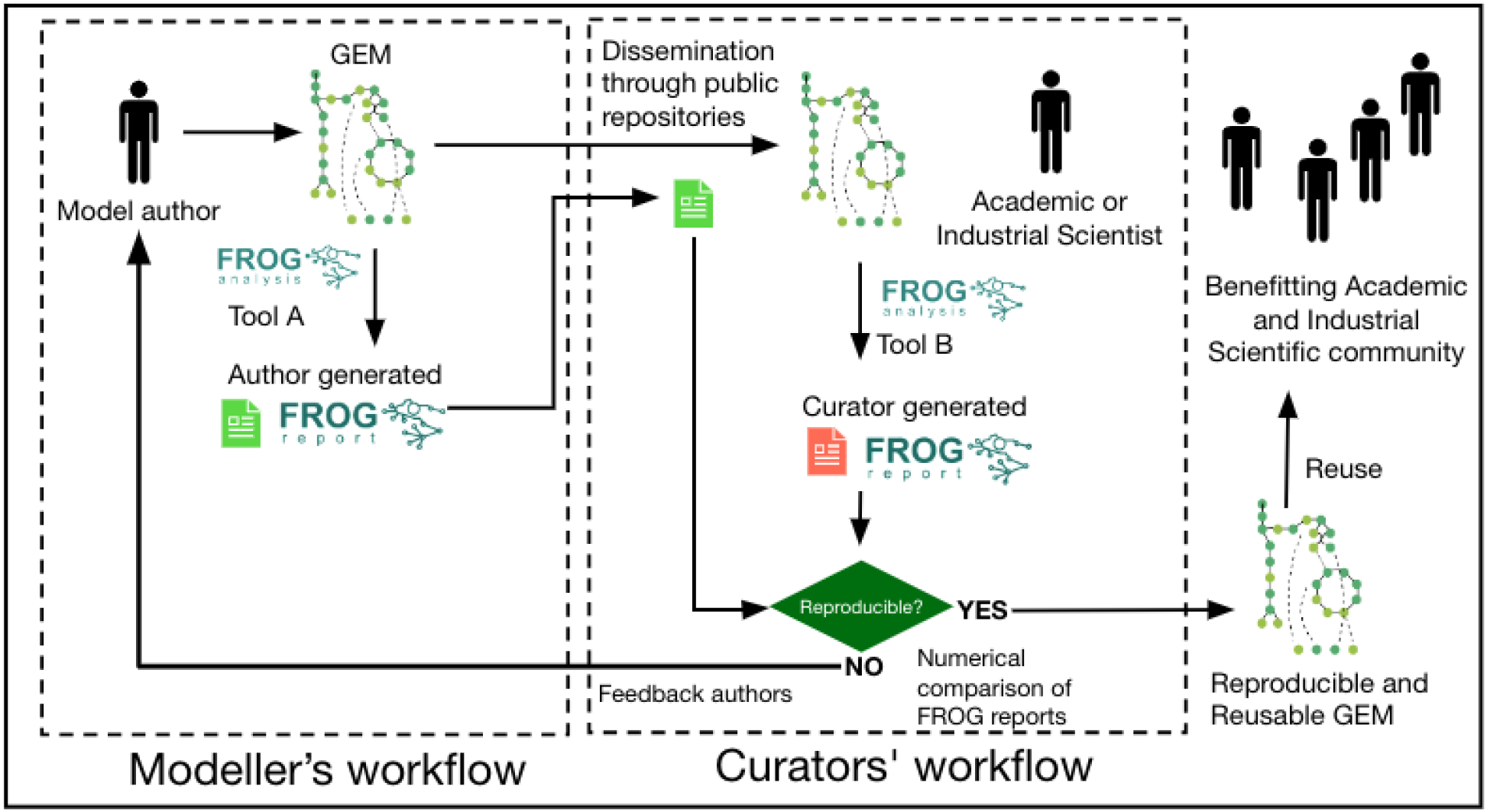
Schematic representation of modellers and curators workflow. Model authors submit GEMs and FROG reports to a public repository. This will allow curators to assess whether the simulations are numerically reproducible using different tools. A public collection of reproducible and reusable GEMs will significantly benefit the wider scientific community.

The FROG community effort generated several open-source tools based on major GEM modelling software, including command-line tools and web interfaces, to run FROG analysis and generate reports (see Table 1). These tools have been harmonised and evaluated to ensure they generate standardised and comparable FROG reports. These publicly shared FROG reports, along with the original models in standard format SBML-FBC (Keating et al., 2020; Olivier and Bergmann, 2018), can now be used by independent modellers, curators or reviewers to autonomously assess the reproducibility of a model by running these standardised analyses in the FROG tools.

**Table 1:**
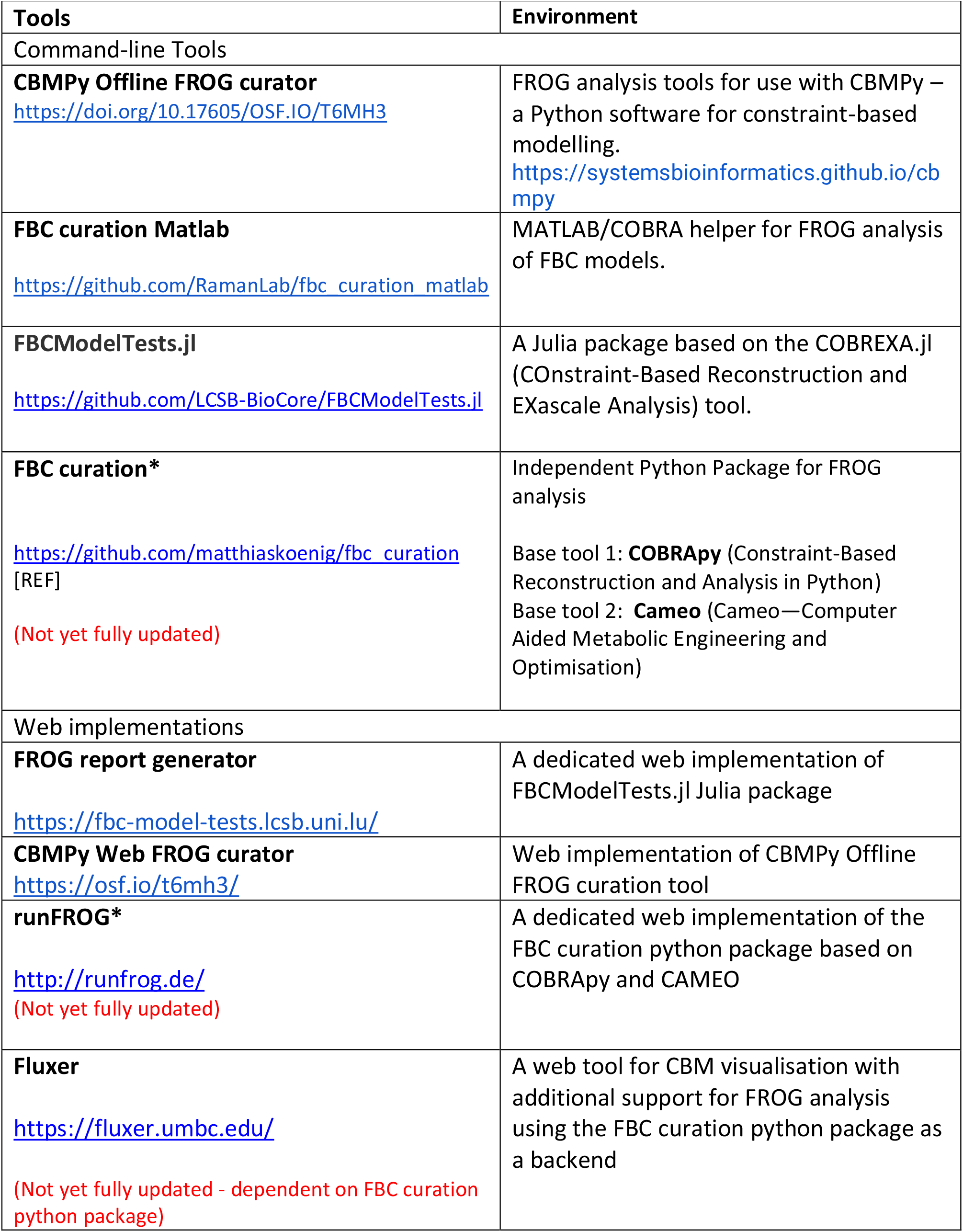
List of FROG tools.

To facilitate a retrospective reproducibility assessment of previously published models and establish a connection between the FROG report and the results presented in manuscripts, we developed a reporting framework to generate a “miniFROG report”. This manually created data table follows a standardised schema, listing results described in the manuscript and corroborating them against the results in the FROG report. Complete specifications for the FROG and miniFROG reports are maintained by the community at https://github.com/EBI-BioModels/frog-specification.

BioModels (Malik-Sheriff et al., 2020), one of the largest repositories of curated biological system models, has integrated FROG analysis into its workflow for curating CBMs and GEMs. To evaluate this approach, BioModels received 65 GEM submissions and their associated FROG reports from the community (see Supplementary Table 1). Out of these 26 models (about 40%) were reproduced without any intervention. For the remaining models, 18 (about 28%) required minor technical interventions for reproduction. In the case of 21 models (around 32%), authors were contacted to either upload the correct version, address SBML validity issues, or resolve other technical problems, such as missing report elements. Ultimately, all models were successfully reproduced, some with a degree of numerical tolerance (see Supplementary Figure 2). FROG reporting allowed the model evaluator to detect issues that hindered the reproduction of the results. These issues include inconsistencies in metadata and data reporting, the validity of SBML, and discrepancies in numerical precision with different constraint solvers. Crucially, the FROG reports allowed rapid identification and communication of such problems to model authors, prospectively enabling a prompt correction to achieve complete reproducibility of the results for the models. Reproduced models are then annotated them with model-level metadata, and generated MEMOTE reports as part of the curation process in BioModels (see Supplementary Table 1).

The standardised FROG analysis, reports, and tools developed by the community and the dedicated model curation in BioModels are crucial in making CBMs and GEMs reproducible and reusable. By providing a reproducibility guarantee for CBMs and GEMs, FROG-based curation will significantly enhance their reuse, extension, and integration into new knowledge generation pipelines, thus fast-forwarding scientific discovery.

## Acknowledgements

The curation was partially carried out using the HPC facilities of the University of Luxembourg (hpc.uni.lu). CBMPy Web was developed on the SURF (www.surf.nl) Research Cloud. P.S. acknowledges the Prime Minister’s Research Fellowship (PMRF) from the Ministry of Education, Government of India. MB was supported by the Systems Biology Grant of the University of Surrey. MK was supported by the Federal Ministry of Education and Research (BMBF, Germany) by grant number 031L0304B and by the German Research Foundation (DFG) by grant number 436883643 and by grant number 465194077. This work was supported by the BMBF-funded de.NBI Cloud within the German Network for Bioinformatics Infrastructure (de.NBI) (031A537B, 031A533A, 031A538A, 031A533B, 031A535A, 031A537C, 031A534A, 031A532B). DYL was supported by the National Research Foundation of Korea (NRF) grants funded by the Korean government (MSIT) (RS-2024-00351458, RS-2024-00341312). MCT was funded by the Portuguese Foundation for Science and Technology (UIDB/04565/2020, UIDP/04565/2020).

## Supplementary Material

### Curation of Constraint-based models (CBMs) / Genome-scale metabolic models (GEMs) in BioModels repository

BioModels is a leading repository of mathematical models of biological systems, hosting over 1050 curated models. Over the past 18 years, the primary focus of curation was ODE-based kinetic models (Malik-Sheriff et al., 2020). Curation in BioModels is a manual process that involves ensuring the model (1) is encoded in a syntactically valid standard format such as SBML (Keating et al., 2020), (2) is reproducible, and (3) is semantically enriched with controlled vocabularies such as GO (The Gene Ontology Consortium, 2019), ChEBI (Hastings et al., 2016), etc. The curation activities ultimately aim at making the models FAIReR (**F**indable, **A**ccessible, **I**nteroperable, **Re**usable, and **R**eproducible), which is extended from the originally suggested FAIR principles (Wilkinson et al., 2016). A model author is expected to submit an SBML model (as the main file), FROG report (as an additional file), and miniFROG (as an additional file) to BioModels. Curators at BioModels will independently try to reproduce the FROG report using a tool different from the one used by the modeller. One of the FROG test suite tools will be used to verify the reproducibility of the CBMs and GEMs submitted. If the results are reproducible, the model will be added to the curated branch of BioModels.

Furthermore, the curator will cross-check the miniFROG report to ensure that the FROG report and the results reported in the manuscript are consistent. Model-level semantic annotations will also be added to the model following MIRIAM guidelines (Le Novère et al., 2005). The quality and consistency of the model annotations will be tested using the MEMOTE test suite, and the MEMOTE report will be uploaded as an additional file. Supplementary Figure 1 summarises the curation process in BioModels—Supplementary Table 1 lists all the models submitted to BioModels with an FROG report. Among the 50 submissions, 24 models were already curated using FROG. These models are reproducible, and model-level annotations were added.

**Supplementary Figure 1:**
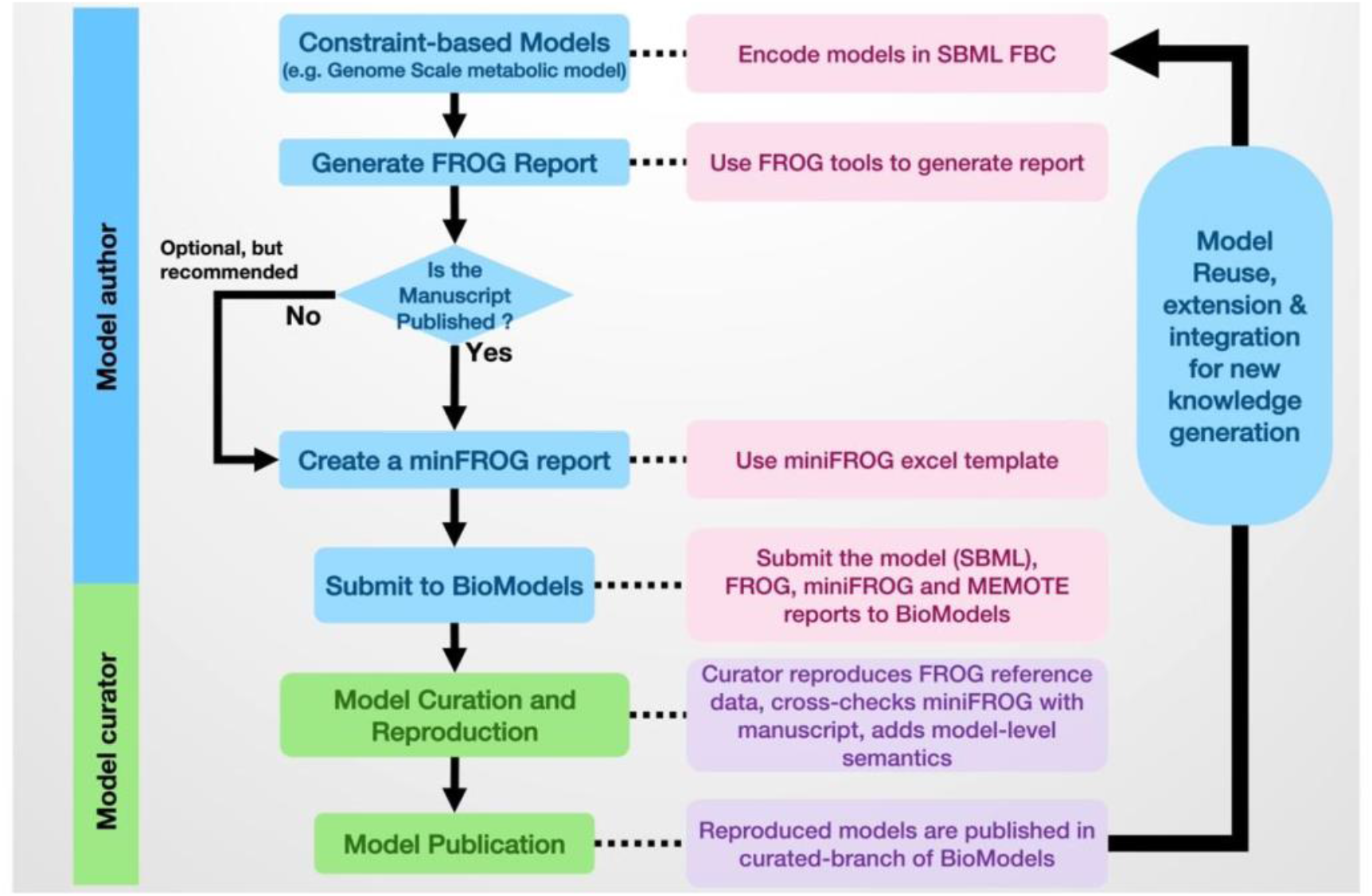
Workflow for the curation of constraint-based models in BioModels using FROG

**Supplementary Figure 2:**
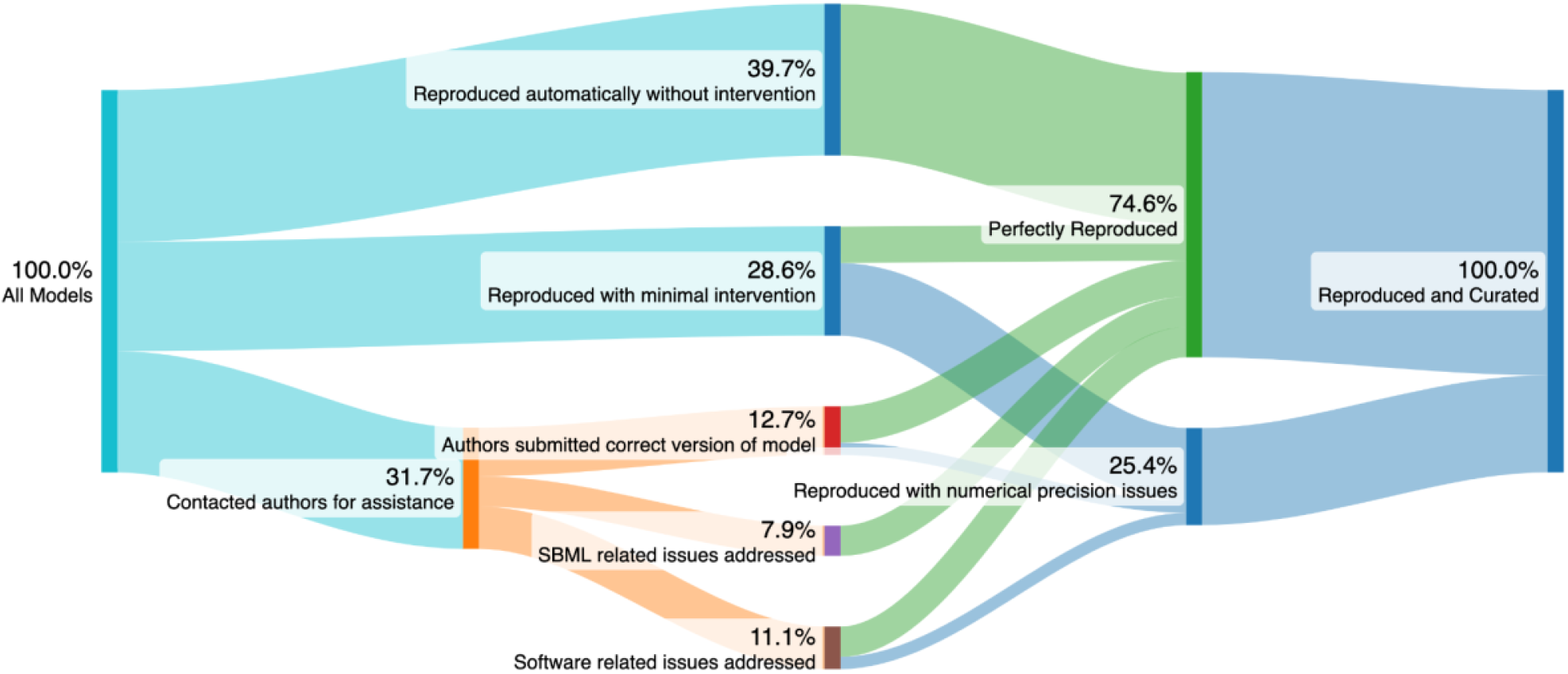
Summary of the models received for the curation and the problems resolved during the curation. Most models were reproducible as-is, either automatically or with minor technical intervention. In approximately one-third of the models, we had to contact authors to upload the correct model version, address SBML validity issues, or fix technical issues such as missing parts of reports. Eventually, the majority of the models were reproduced perfectly, and around a quarter was reproduced with controllable numeric precision issues caused by floating-point round-off inflation, with appropriate notices added for the users.

**Supplementary Table 1:**
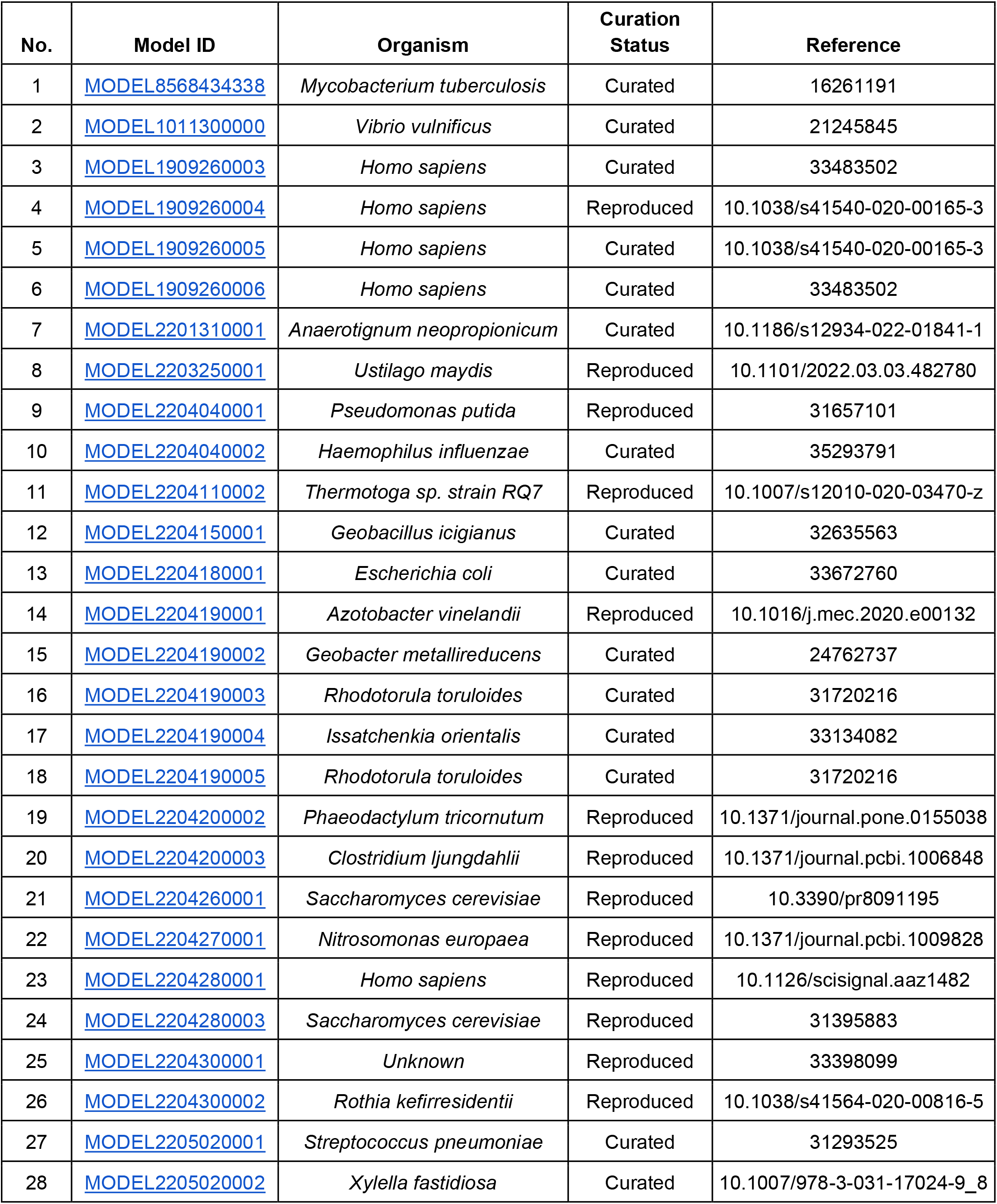
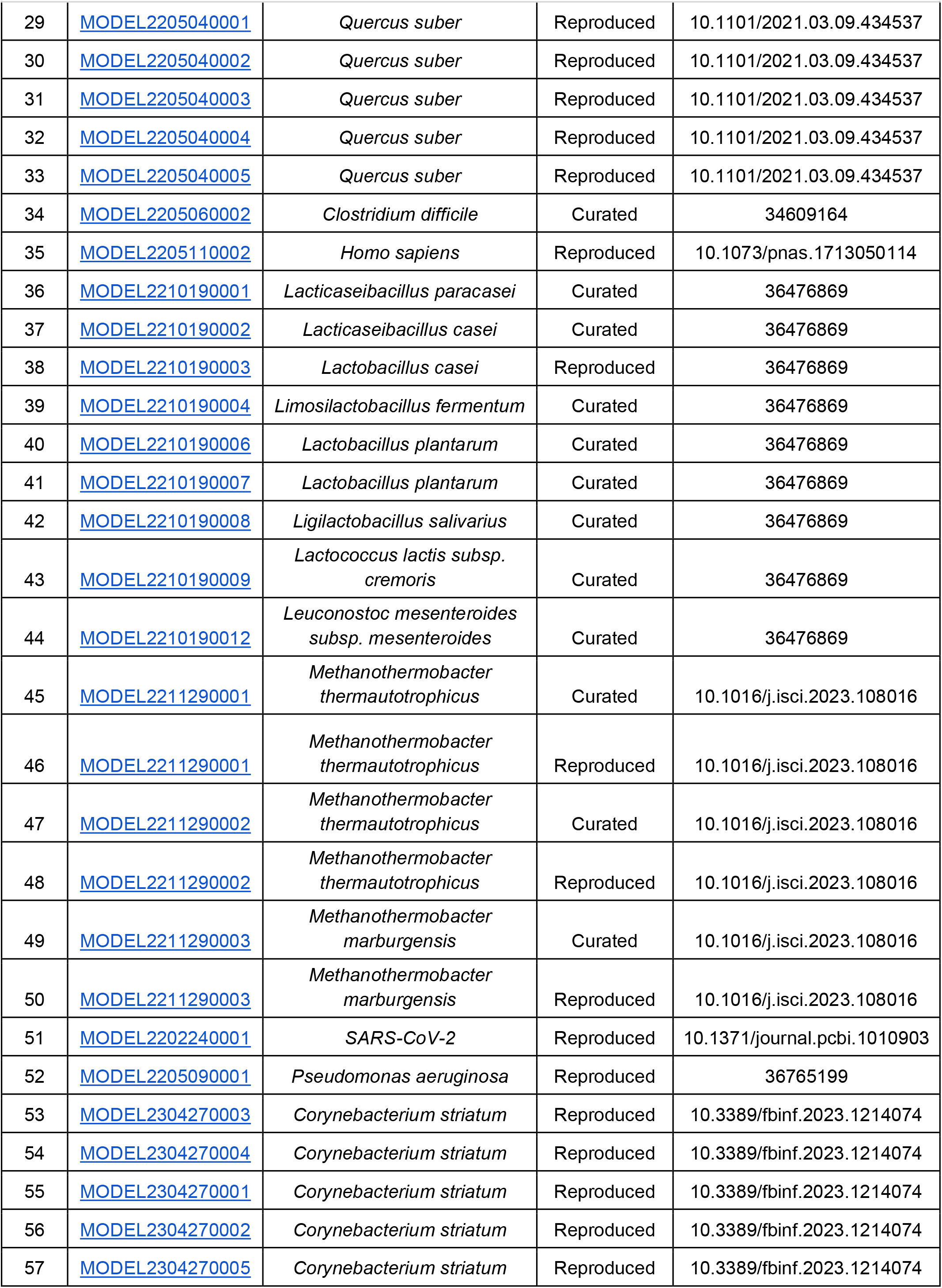
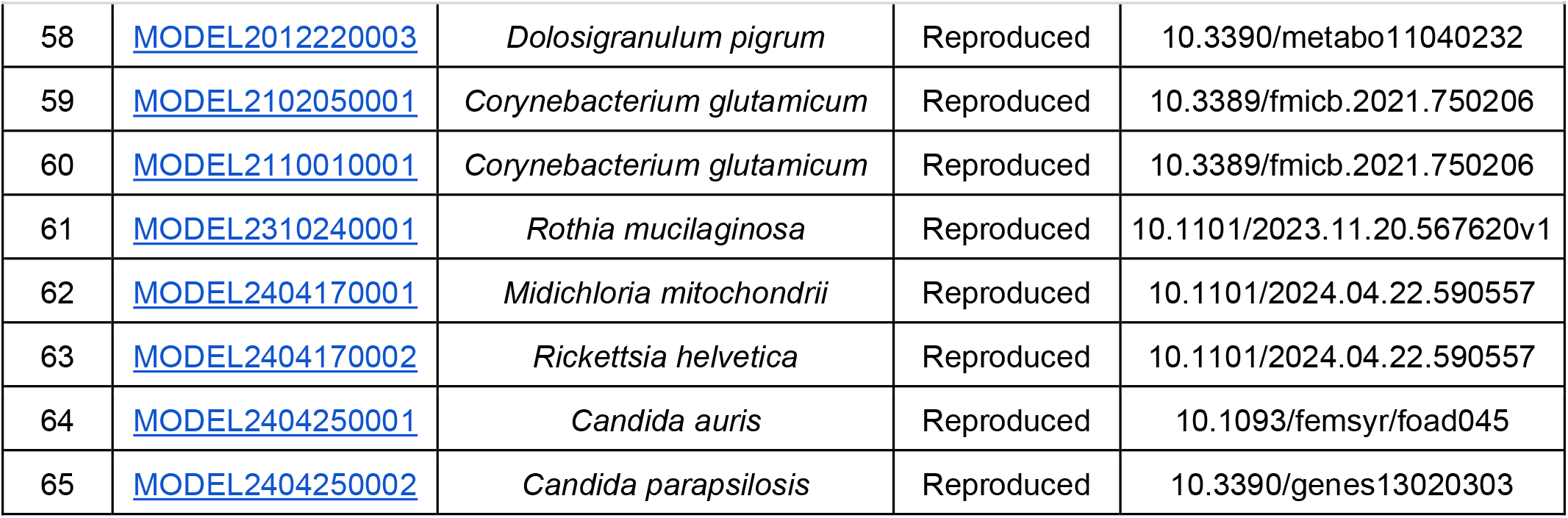
List of GEMs submitted to BioModels for FROG-based curation. Curated models are reproduced and annotated with model-level metadata cross-referencing appropriate ontology terms and submitted to BioModels. Reproduced models are awaiting model annotation.

| No. | Model ID | Organism | Curation Status | Reference |
| --- | --- | --- | --- | --- |
| 1 | MODEL8568434338 | *Mycobacterium tuberculosis* | Curated | 16261191 |
| 2 | MODEL1011300000 | *Vibrio vulnificus* | Curated | 21245845 |
| 3 | MODEL1909260003 | *Homo sapiens* | Curated | 33483502 |
| 4 | MODEL1909260004 | *Homo sapiens* | Reproduced | 10.1038/s41540-020-00165-3 |
| 5 | MODEL1909260005 | *Homo sapiens* | Curated | 10.1038/s41540-020-00165-3 |
| 6 | MODEL1909260006 | *Homo sapiens* | Curated | 33483502 |
| 7 | MODEL2201310001 | *Anaerotignum neopropionicum* | Curated | 10.1186/s12934-022-01841-1 |
| 8 | MODEL2203250001 | *Ustilago maydis* | Reproduced | 10.1101/2022.03.03.482780 |
| 9 | MODEL2204040001 | *Pseudomonas putida* | Reproduced | 31657101 |
| 10 | MODEL2204040002 | *Haemophilus influenzae* | Curated | 35293791 |
| 11 | MODEL2204110002 | *Thermotoga sp. strain RQ7* | Reproduced | 10.1007/s12010-020-03470-z |
| 12 | MODEL2204150001 | *Geobacillus icigianus* | Curated | 32635563 |
| 13 | MODEL2204180001 | *Escherichia coli* | Curated | 33672760 |
| 14 | MODEL2204190001 | *Azotobacter vinelandii* | Reproduced | 10.1016/j.mec.2020.e00132 |
| 15 | MODEL2204190002 | *Geobacter metallireducens* | Curated | 24762737 |
| 16 | MODEL2204190003 | *Rhodotorula toruloides* | Curated | 31720216 |
| 17 | MODEL2204190004 | *Issatchenkia orientalis* | Curated | 33134082 |
| 18 | MODEL2204190005 | *Rhodotorula toruloides* | Curated | 31720216 |
| 19 | MODEL2204200002 | *Phaeodactylum tricornutum* | Reproduced | 10.1371/journal.pone.0155038 |
| 20 | MODEL2204200003 | *Clostridium ljungdahlii* | Reproduced | 10.1371/journal.pcbi.1006848 |
| 21 | MODEL2204260001 | *Saccharomyces cerevisiae* | Reproduced | 10.3390/pr8091195 |
| 22 | MODEL2204270001 | *Nitrosomonas europaea* | Reproduced | 10.1371/journal.pcbi.1009828 |
| 23 | MODEL2204280001 | *Homo sapiens* | Reproduced | 10.1126/scisignal.aaz1482 |
| 24 | MODEL2204280003 | *Saccharomyces cerevisiae* | Reproduced | 31395883 |
| 25 | MODEL2204300001 | Unknown | Reproduced | 33398099 |
| 26 | MODEL2204300002 | *Rothia kefirresidentii* | Reproduced | 10.1038/s41564-020-00816-5 |
| 27 | MODEL2205020001 | *Streptococcus pneumoniae* | Curated | 31293525 |
| 28 | MODEL2205020002 | *Xylella fastidiosa* | Curated | 10.1007/978-3-031-17024-9_8 |
| FROG Analysis |  |  |  |  |
| 29 | MODEL2205040001 | *Quercus suber* | Reproduced | 10.1101/2021.03.09.434537 |
| 30 | MODEL2205040002 | *Quercus suber* | Reproduced | 10.1101/2021.03.09.434537 |
| 31 | MODEL2205040003 | *Quercus suber* | Reproduced | 10.1101/2021.03.09.434537 |
| 32 | MODEL2205040004 | *Quercus suber* | Reproduced | 10.1101/2021.03.09.434537 |
| 33 | MODEL2205040005 | *Quercus suber* | Reproduced | 10.1101/2021.03.09.434537 |
| 34 | MODEL2205060002 | *Clostridium difficile* | Curated | 34609164 |
| 35 | MODEL2205110002 | *Homo sapiens* | Reproduced | 10.1073/pnas.1713050114 |
| 36 | MODEL2210190001 | *Lactocaseibacillus paracasei* | Curated | 36476869 |
| 37 | MODEL2210190002 | *Lactocaseibacillus casei* | Curated | 36476869 |
| 38 | MODEL2210190003 | *Lactobacillus casei* | Reproduced | 36476869 |
| 39 | MODEL2210190004 | *Limosilactobacillus fermentum* | Curated | 36476869 |
| 40 | MODEL2210190006 | *Lactobacillus plantarum* | Curated | 36476869 |
| 41 | MODEL2210190007 | *Lactobacillus plantarum* | Curated | 36476869 |
| 42 | MODEL2210190008 | *Ligilactobacillus salivarius* | Curated | 36476869 |
| 43 | MODEL2210190009 | *Lactococcus lactis* subsp.<br>*cremoris* | Curated | 36476869 |
| 44 | MODEL2210190012 | *Leuconostoc mesenteroides*<br>subsp. *mesenteroides* | Curated | 36476869 |
| 45 | MODEL2211290001 | *Methanothermobacter*<br>*thermautotrophicus* | Curated | 10.1016/j.isci.2023.108016 |
| 46 | MODEL2211290001 | *Methanothermobacter*<br>*thermautotrophicus* | Reproduced | 10.1016/j.isci.2023.108016 |
| 47 | MODEL2211290002 | *Methanothermobacter*<br>*thermautotrophicus* | Curated | 10.1016/j.isci.2023.108016 |
| 48 | MODEL2211290002 | *Methanothermobacter*<br>*thermautotrophicus* | Reproduced | 10.1016/j.isci.2023.108016 |
| 49 | MODEL2211290003 | *Methanothermobacter*<br>*marburgensis* | Curated | 10.1016/j.isci.2023.108016 |
| 50 | MODEL2211290003 | *Methanothermobacter*<br>*marburgensis* | Reproduced | 10.1016/j.isci.2023.108016 |
| 51 | MODEL2202240001 | SARS-CoV-2 | Reproduced | 10.1371/journal.pcbi.1010903 |
| 52 | MODEL2205090001 | *Pseudomonas aeruginosa* | Reproduced | 36765199 |
| 53 | MODEL2304270003 | *Corynebacterium striatum* | Reproduced | 10.3389/fbinf.2023.1214074 |
| 54 | MODEL2304270004 | *Corynebacterium striatum* | Reproduced | 10.3389/fbinf.2023.1214074 |
| 55 | MODEL2304270001 | *Corynebacterium striatum* | Reproduced | 10.3389/fbinf.2023.1214074 |
| 56 | MODEL2304270002 | *Corynebacterium striatum* | Reproduced | 10.3389/fbinf.2023.1214074 |
| 57 | MODEL2304270005 | *Corynebacterium striatum* | Reproduced | 10.3389/fbinf.2023.1214074 |
| 58 | MODEL2012220003 | *Dolosigranulum pigrum* | Reproduced | 10.3390/metabo11040232 |
| 59 | MODEL2102050001 | *Corynebacterium glutamicum* | Reproduced | 10.3389/fmicb.2021.750206 |
| 60 | MODEL2110010001 | *Corynebacterium glutamicum* | Reproduced | 10.3389/fmicb.2021.750206 |
| 61 | MODEL2310240001 | *Rothia mucilaginosa* | Reproduced | 10.1101/2023.11.20.567620v1 |
| 62 | MODEL2404170001 | *Midichloria mitochondrii* | Reproduced | 10.1101/2024.04.22.590557 |
| 63 | MODEL2404170002 | *Rickettsia helvetica* | Reproduced | 10.1101/2024.04.22.590557 |
| 64 | MODEL2404250001 | *Candida auris* | Reproduced | 10.1093/femsyr/foad045 |
| 65 | MODEL2404250002 | *Candida parapsilosis* | Reproduced | 10.3390/genes13020303 |

## References

Anton, M., Almaas, E., Benfeitas, R., Benito-Vaquerizo, S., Blank, L.M., Dräger, A., Hancock, J.M., Kittikunapong, C., König, M., Li, F., Liebal, U.W., Lu, H., Ma, H., Mahadevan, R., Mardinoglu, A., Nielsen, J., Nogales, J., Pagni, M., Papin, J.A., Patil, K.R., Price, N.D., Robinson, J.L., Sánchez, B.J., Suarez-Diez, M., Sulheim, S., Svensson, L.T., Teusink, B., Vongsangnak, W., Wang, H., Zeidan, A.A., Kerkhoven, E.J., 2023. standard-GEM: standardization of open-source genome-scale metabolic models. 10.1101/2023.03.21.512712

The Gene Ontology Consortium, 2019. The Gene Ontology Resource: 20 years and still GOing strong. Nucleic Acids Res. 47, D330–D338. 10.1093/nar/gky1055

Gu, C., Kim, G.B., Kim, W.J., Kim, H.U., Lee, S.Y., 2019. Current status and applications of genome-scale metabolic models. Genome Biol 20, 121. 10.1186/s13059-019-1730-3

Hastings, J., Owen, G., Dekker, A., Ennis, M., Kale, N., Muthukrishnan, V., Turner, S., Swainston, N., Mendes, P., Steinbeck, C., 2016. ChEBI in 2016: Improved services and an expanding collection of metabolites. Nucleic Acids Res. 44, D1214–1219. 10.1093/nar/gkv1031

Ibrahim, M., Raajaraam, L., Raman, K., 2021. Modelling microbial communities: Harnessing consortia for biotechnological applications. Comput Struct Biotechnol J 19, 3892–3907. 10.1016/j.csbj.2021.06.048

Keating, S.M., Waltemath, D., König, M., Zhang, F., Dräger, A., Chaouiya, C., Bergmann, F.T., Finney, A., Gillespie, C.S., Helikar, T., Hoops, S., Malik-Sheriff, R.S., Moodie, S.L., Moraru, I.I., Myers, C.J., Naldi, A., Olivier, B.G., Sahle, S., Schaff, J.C., Smith, L.P., Swat, M.J., Thieffry, D., Watanabe, L., Wilkinson, D.J., Blinov, M.L., Begley, K., Faeder, J.R., Gómez, H.F., Hamm, T.M., Inagaki, Y., Liebermeister, W., Lister, A.L., Lucio, D., Mjolsness, E., Proctor, C.J., Raman, K., Rodriguez, N., Shaffer, C.A., Shapiro, B.E., Stelling, J., Swainston, N., Tanimura, N., Wagner, J., Meier-Schellersheim, M., Sauro, H.M., Palsson, B., Bolouri, H., Kitano, H., Funahashi, A., Hermjakob, H., Doyle, J.C., Hucka, M., SBML Level 3 Community members, 2020. SBML Level 3: an extensible format for the exchange and reuse of biological models. Molecular Systems Biology 16, e9110. 10.15252/msb.20199110

Le Novère, N., Finney, A., Hucka, M., Bhalla, U.S., Campagne, F., Collado-Vides, J., Crampin, E.J., Halstead, M., Klipp, E., Mendes, P., Nielsen, P., Sauro, H., Shapiro, B., Snoep, J.L., Spence, H.D., Wanner, B.L., 2005. Minimum information requested in the annotation of biochemical models (MIRIAM). Nat. Biotechnol. 23, 1509–1515.

Li, F., Chen, Y., Gustafsson, J., Wang, H., Wang, Y., Zhang, C., Xing, X., 2023. Genome-scale metabolic models applied for human health and biopharmaceutical engineering. Quantitative Biology 11, 363–375. 10.1002/qub2.21

Lieven, C., Beber, M.E., Olivier, B.G., Bergmann, F.T., Ataman, M., Babaei, P., Bartell, J.A., Blank, L.M., Chauhan, S., Correia, K., Diener, C., Dräger, A., Ebert, B.E., Edirisinghe, J.N., Faria, J.P., Feist, A.M., Fengos, G., Fleming, R.M.T., García-Jiménez, B., Hatzimanikatis, V., van Helvoirt, W., Henry, C.S., Hermjakob, H., Herrgård, M.J., Kaafarani, A., Kim, H.U., King, Z., Klamt, S., Klipp, E., Koehorst, J.J., König, M., Lakshmanan, M., Lee, D.-Y., Lee, S.Y., Lee, S., Lewis, N.E., Liu, F., Ma, H., Machado, D., Mahadevan, R., Maia, P., Mardinoglu, A., Medlock, G.L., Monk, J.M., Nielsen, J., Nielsen, L.K., Nogales, J., Nookaew, I., Palsson, B.O., Papin, J.A., Patil, K.R., Poolman, M., Price, N.D., Resendis-Antonio, O., Richelle, A., Rocha, I., Sánchez, B.J., Schaap, P.J., Malik Sheriff, R.S., Shoaie, S., Sonnenschein, N., Teusink, B., Vilaça, P., Vik, J.O., Wodke, J.A.H., Xavier, J.C., Yuan, Q., Zakhartsev, M., Zhang, C., 2020. MEMOTE for standardized genome-scale metabolic model testing. Nature Biotechnology 38, 272–276. 10.1038/s41587-020-0446-y

Malik-Sheriff, R.S., Glont, M., Nguyen, T.V.N., Tiwari, K., Roberts, M.G., Xavier, A., Vu, M.T., Men, J., Maire, M., Kananathan, S., Fairbanks, E.L., Meyer, J.P., Arankalle, C., Varusai, T.M., Knight-Schrijver, V., Li, L., Dueñas-Roca, C., Dass, G., Keating, S.M., Park, Y.M., Buso, N., Rodriguez, N., Hucka, M., Hermjakob, H., 2020. BioModels-15 years of sharing computational models in life science. Nucleic Acids Res. 48, D407–D415. 10.1093/nar/gkz1055

McCloskey, D., Palsson, B.Ø., Feist, A.M., 2013. Basic and applied uses of genome-scale metabolic network reconstructions of Escherichia coli. Mol Syst Biol 9, 661. 10.1038/msb.2013.18

Oberhardt, M.A., Palsson, B.Ø., Papin, J.A., 2009. Applications of genome-scale metabolic reconstructions. Mol Syst Biol 5, 320. 10.1038/msb.2009.77

Olivier, B.G., Bergmann, F.T., 2018. SBML Level 3 Package: Flux Balance Constraints version 2. J Integr Bioinform 15, 20170082. 10.1515/jib-2017-0082

Ravikrishnan, A., Raman, K., 2015. Critical assessment of genome-scale metabolic networks: the need for a unified standard. Briefings in Bioinformatics 16, 1057–1068. 10.1093/bib/bbv003

Renz, A., Widerspick, L., Dräger, A., 2020. FBA reveals guanylate kinase as a potential target for antiviral therapies against SARS-CoV-2. Bioinformatics 36, i813–i821. 10.1093/bioinformatics/btaa813

Schellenberger, J., Que, R., Fleming, R.M.T., Thiele, I., Orth, J.D., Feist, A.M., Zielinski, D.C., Bordbar, A., Lewis, N.E., Rahmanian, S., Kang, J., Hyduke, D.R., Palsson, B.Ø., 2011. Quantitative prediction of cellular metabolism with constraint-based models: the COBRA Toolbox v2.0. Nat Protoc 6, 1290–1307. 10.1038/nprot.2011.308

Tiwari, K., Kananathan, S., Roberts, M.G., Meyer, J.P., Sharif Shohan, M.U., Xavier, A., Maire, M., Zyoud, A., Men, J., Ng, S., Nguyen, T.V.N., Glont, M., Hermjakob, H., Malik-Sheriff, R.S., 2021. Reproducibility in systems biology modelling. Molecular Systems Biology 17, e9982. 10.15252/msb.20209982

van der Ark, K.C.H., van Heck, R.G.A., Martins Dos Santos, V.A.P., Belzer, C., de Vos, W.M., 2017. More than just a gut feeling: constraint-based genome-scale metabolic models for predicting functions of human intestinal microbes. Microbiome 5, 78. 10.1186/s40168-017-0299-x

Wilkinson, M.D., Dumontier, M., Aalbersberg, Ij.J., Appleton, G., Axton, M., Baak, A., Blomberg, N., Boiten, J.-W., da Silva Santos, L.B., Bourne, P.E., Bouwman, J., Brookes, A.J., Clark, T., Crosas, M., Dillo, I., Dumon, O., Edmunds, S., Evelo, C.T., Finkers, R., Gonzalez-Beltran, A., Gray, A.J.G., Groth, P., Goble, C., Grethe, J.S., Heringa, J., ‘t Hoen, P.A.C., Hooft, R., Kuhn, T., Kok, R., Kok, J., Lusher, S.J., Martone, M.E., Mons, A., Packer, A.L., Persson, B., Rocca-Serra, P., Roos, M., van Schaik, R., Sansone, S.-A., Schultes, E., Sengstag, T., Slater, T., Strawn, G., Swertz, M.A., Thompson, M., van der Lei, J., van Mulligen, E., Velterop, J., Waagmeester, A., Wittenburg, P., Wolstencroft, K., Zhao, J., Mons, B., 2016. The FAIR Guiding Principles for scientific data management and stewardship. Sci Data 3. 10.1038/sdata.2016.18

